# Biological Evaluation of Molecules of the azaBINOL Class as Antiviral Agents: Specific Inhibition of HIV-1 RNase H Activity by 7-Isopropoxy-8-(naphth-1-yl)quinoline

**DOI:** 10.1101/525105

**Authors:** Ross D. Overacker, Somdev Banerjee, George F. Neuhaus, Selena Milicevic Sephton, Alexander Herrmann, James A. Strother, Ruth Brack-Werner, Paul R. Blakemore, Sandra Loesgen

## Abstract

Inspired by bioactive biaryl-containing natural products found in plants and the marine environment, a series of synthetic compounds belonging to the azaBINOL chiral ligand family was evaluated for antiviral activity against HIV-1. Testing of 39 unique azaBINOLs in a single-round infectivity assay resulted in the identification of three promising antiviral compounds, including 7-isopropoxy-8-(naphth-1-yl)quinoline (azaBINOL **B#24**), which exhibited low-micromolar activity. The active compounds and several close structural analogues were further tested against three different HIV-1 envelope pseudotyped viruses as well as in a full-virus replication system (EASY-HIT). Mode-of-action studies using a time-of-addition assay indicated that azaBINOL **B#24** acts after viral entry but before viral assembly and budding. HIV-1 reverse transcriptase (RT) assays that individually test for polymerase and RNase H activity were used to demonstrate that **B#24** inhibits RNase H activity, most likely allosterically. Further binding analysis using bio-layer interferometry (BLI) showed that **B#24** interacts with HIV-1 RT in a highly specific manner. These results indicate that azaBINOL **B#24** is a potentially viable, novel lead for the development of new HIV-1 RNase H inhibitors. Furthermore, this study demonstrates that the survey of libraries of synthetic compounds, designed purely with the goal of facilitating chemical synthesis in mind, may yield unexpected and selective drug leads for the development of new antiviral agents.

## 1. Introduction

HIV/AIDS continues to be a major global health epidemic. In 2017, there were 36.9 million people living with HIV worldwide and an additional 1.8 million people became newly infected.^1^ Despite substantial efforts towards vaccine development, there are currently no FDA-approved vaccines and management of HIV infection requires long-term treatment with potent anti-HIV drugs.^2^ Although drug regimens such as antiretroviral therapy (ART) are able to keep HIV viral load low in infected patients, these treatments are typically limited by adverse side effects, increasing drug resistance, high costs, and global availability shortages.^3^ Currently, therapeutic antiviral drugs target different phases of the HIV lifecycle including viral attachment, fusion, reverse transcription, integration, and protease activity. Treatments often utilize multiple drugs in combination to combat the rapid emergence of chemoresistant viruses.^3^ Therefore, novel therapeutics that act on previously untargeted steps of the viral life cycle are urgently needed to circumvent the onset of drug resistance and to improve treatments.^4^

HIV reverse transcriptase (RT) has been successfully targeted with first and second-generation non-nucleosides reverse transcriptase inhibitors (NNRTIs) as exemplified by nevirapine, first introduced in 1996, and more recently rilpivirine, in 2011.^5, 6^ Out of the 27 currently FDA approved HIV drugs on the market, 13 of them target RT polymerase activity, including the NNRTIs.^7^ However, the reverse transcriptase enzyme is a multifunctional protein and no drugs have been developed yet that target the RT ribonuclease H (RNase H) activity, which has recently been validated as a target for small molecule drug intervention.^8, 9^ HIV RT is responsible for converting the single-stranded viral RNA genome to a double-stranded DNA for subsequent integration into the genome of the host cell. The heterodimeric RT protein (p66/p51) has separate active sites for polymerase and RNase H activity. The polymerase starts the synthesis of DNA by first copying the viral RNA genome and forming RNA:DNA hybrids while RNase H catalyzes the degradation of RNA in DNA:RNA hybrids to finally form duplex DNA.^10^ The RNase H active site contains two bivalent Mg^2+^ ions that chelate and cleave the RNA phosphate backbone by directing a nucleophilic water molecule towards the phosphate linkage.^11^ It has been shown that RNase H activity can be abolished through Mg^2+^ chelation in the active site or through allosteric binding near the NNRTI site causing conformational changes.^9^ The addition of novel antiviral drugs that target RNase H activity to current combinatorial regimens would introduce a new synergistic method of HIV-1 inhibition greatly improving efficacy of treatment options.

To identify novel leads for drug discovery efforts one may look to sources of compounds that have been either infrequent explored or else untapped in prior studies. Notable in this regard are the numerous ostensibly artificial organic molecules that have been introduced as chiral metal ligands and/or organocatalysts for the purpose of facilitating catalytic enantioselective syntheses. The structural features present in such molecules that are necessary for their intended function (e.g., chiral scaffolds with few rotatable bonds, donor sites from atoms with lone pairs, hydrogen-bond acceptors and donors, sites of localized charge density, zones of steric encumbrance, etc.) could also lead to meaningful and potentially specific interactions with proteins and other classes of biomolecules involved in various diseases. Axially chiral biaryl compounds based on 1,1′-binaphthyl scaffolds are considered a ‘privileged’ class of reagents for enantioselective synthesis and the principal member of this group, 1,1′-bi-2-naphthol (BINOL, 1), has become one of the most widely used ligands for stoichiometric and catalytic asymmetric reactions.^12^ While BINOL itself has previously been found to be cytotoxic, many other biaryl compounds, either found in nature or of artificial origin, have shown potent and selective bioactivities.^13,14^ For example, the axially chiral dimeric naphthylisoquinolone alkaloids first isolated from *Anicistrocladus korupensis* in 1991 and later named the michellamines, exhibit selective anti-HIV activity.^15–19^ Given these facts taken together with the existence of other antiviral biaryl natural products (e.g., dioncophylline^20^) and recently identified synthetic antiviral drug leads with multiple aromatic ring systems (e.g., arbidol,^21^ peptide triazoles,^22^ rhodanine derivatives,^23^ naphthylhydrazones,^24^ and hydroxypyridones^25^), we elected to test a library of heterocyclic biaryl compounds available to us and belonging to the so-called ‘azaBINOL’ chiral ligand family for inhibition of HIV-1 infection.

The azaBINOLs are nitrogenous analogs of BINOL based on isostructural 8-(naphth-1-yl)quinoline (**2**, 8-azaBINOL)^26^ and 8,8’-biquinolyl (3, 8,8′-diazaBINOL)^27–29^ motifs (Figure 1). These molecules have been a focus of interest both from a fundamental standpoint^26, 30, 31^ and for their potential utility in enantioselective synthesis,^32, 33^ but prior to this work, studies of any aspect of the biological activity of azaBINOLs had yet to be reported. Herein, we show that deoxy-8-azaBINOL derivatives provide a novel scaffold for the inhibition of HIV-1 RT RNase H activity. Our lead compound, the isopropyl ether derivative of 2’-deoxy-8-azaBINOL (**B#24**), shows unoptimized low micromolar activity (4-9 μM range) in an HIV single round infectivity assay as well as in fully infectious viral assays with low cytotoxicity and a selectivity index of 14.

**Figure 1.**
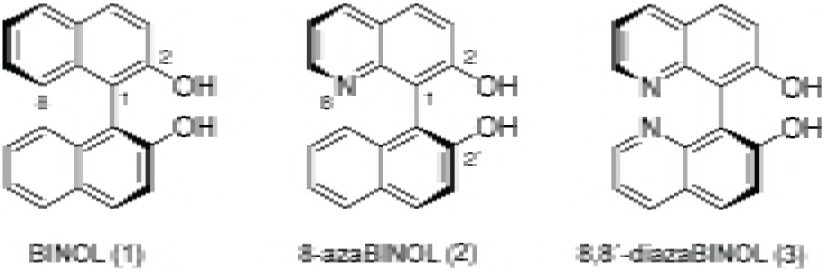
1,1′-Bi-2-naphthol (BINOL, **1**) and related 8-(naphth-1-yl)quinoline (**2**) and 8,8’;-biquinolyl (3) compounds belonging to the ‘azaBINOL’ class. The colloquial names ‘8-azaBINOL’ and ‘8,8′-diazaBINOL’ conform to BINOL atom numbering. Molecules are depicted in (*aS*)-configuration and can exist in racemic or scalemic form.

## 2. Chemistry

A significant advantage of the azaBINOL family of molecules as compared to their all carbocyclic BINOL congeners is the ease of derivatization of the quinoline nucleus and therefore the facility with which essentially any position of azaBINOL scaffolds can be decorated with ancilliary functionality.^29^ Of the six compounds of primary interest herein (*vide infra*, see Figure 2), only the quinol-type 2′-deoxy-8-azaBINOL carbamate derivative **B#43** was previously described in the literature.^26^ The five new compounds were prepared from known deoxy azaBINOLs (4, 5, and 6) via straightforward alkylation and acylation reactions (Scheme 1). Carbamate **B#43**, itself accessed by Suzuki-Miyauri cross-coupling of an 8-iodoquinoline and 1-naphthaleneboronic acid,^26^ was converted to isopropyl ether **B#24** by saponification to quinol 4 followed by Williamson ether synthesis (Scheme 1). The other four compounds, naphthol-type 2-deoxy-8-azaBINOL derivatives **B#59** and **B#60** and 2-deoxy-8,8′-diazaBINOL derivatives **B#57** and **B#58**, were prepared from the corresponding phenols **5** and **6**. As hitherto reported, phenols 5 and 6 are themselves efficiently prepared by N-directed oxidative CH functionalization of 8-(naphth-1-yl)quinoline^26^ and 8,8′-biquinolyl,^29^ respectively. All six of the azaBINOLs of main focus were prepared and tested for biological activity in racemic form. The configurational stability of these axially chiral compounds has yet to be determined, however, their racemization half-lives are likely to be significantly higher than those of quinol **4** [τ_1/2(rac.)_ = 120 h in MeOH at 24 °C] and naphthol **5** [τ_1/2(rac.)_ = 89 h in MeOH at 24 °C] which have been measured as indicated.^26^

**Figure 2.**
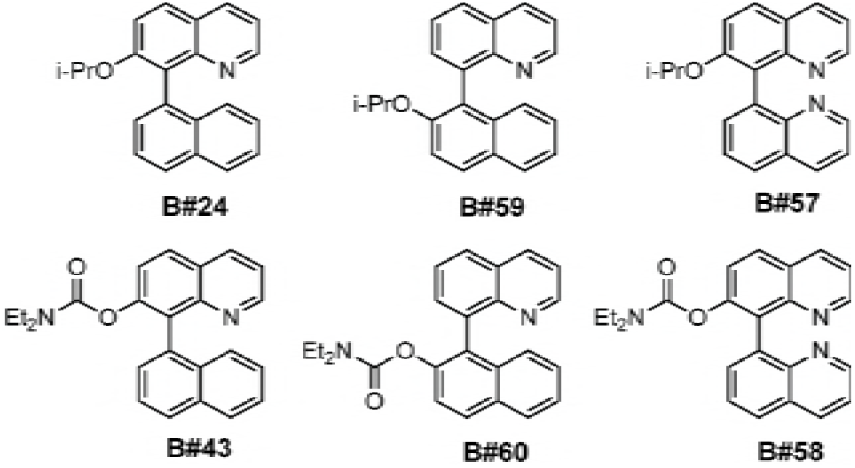
Structures of selected azaBINOL compounds from synthetic library that show significant antiviral activity at single dose concentrations (**B#24**, **B#43**, **B#60**) along with closely related congeners showing only modest antiviral activity (**B#59**, **B#58**, **B#57**).

**Scheme 1.**
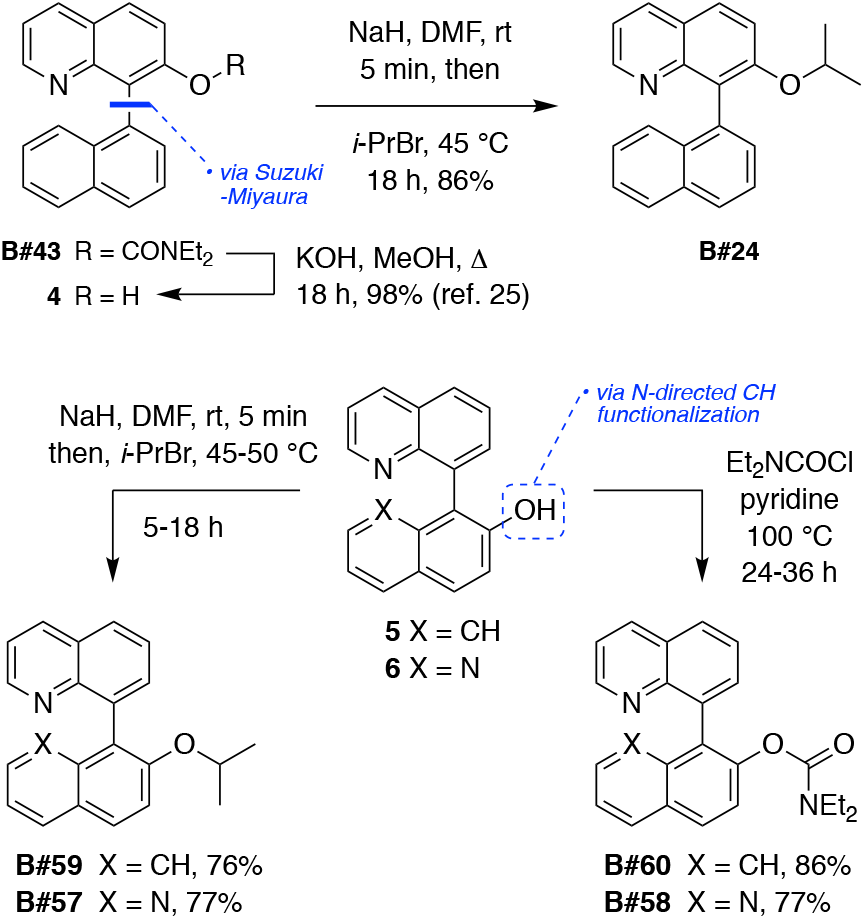
Synthesis of deoxy-8-azaBINOL (**B#24, B#43, B#59**, and **B#60**) and 2-deoxy-8,8′-diazaBINOL (**B#57** and **B#58**) derivatives of primary interest for antiviral activity evaluation.

## 3. Results

### 3.1. HIV-1 ***in vitro*** activity screening of azaBINOL compounds

A library of 39 unique azaBINOLs and two BINOLs was screened for antiviral HIV-1 activity using a pseudo-typed viral particle, single round infectivity assay (HIVpp) (see Supporting Information for full screening data and structures of all library members; all compounds were screened in racemic form and four were additionally evaluated as their enantiopure (*aS*)- and (*aR*)-atropisomers). Antiviral activity was compared directly to compound cytotoxicity using a standard MTT-based cell viability assay to assess selectivity indices of compounds. Initial screening efforts at single-dose concentrations (10 μg/mL) of the full compound library revealed three compounds with low micromolar anti-HIV activity: **B#24**, **B#43**, and **B#60** (Figure 2). These compounds, an isopropyl ether (**B#24**) and two carbamate derivatives (**B#43** and **B#60**) of deoxy-8-azaBINOL molecules, showed high viral inhibitory activity with only minor cytotoxicity and warranted further investigation. Closer analysis of the compound library revealed three additional azaBINOLs sharing similar structural features to the aforementioned active compounds: a naphthol-type regioisomer of quinol-type ether **B#24** (**B#59**), as well as 2-deoxy-8,8′-diazaBINOL congeners of the isopropyl ethers and carbamates (**B#57** and **B#58**). Despite the close structural similarities of these compounds, they showed little antiviral activity as compared to **B#24**, **B#43**, and **B#60** (Figure 2). HIV antiviral activity was evident for the isopropyl ether and carbamate derivatives of 2- and 2′-deoxy-8-azaBINOL but it was notably completely absent for the corresponding 2-deoxy-8,8′-diazaBINOL series of compounds. Due to the close structural similarities between the active isopropyl ether and carbamate compounds to their non-active counterparts in the HIVpp assay, we decided to move all six compounds forward for further activity explorations.

Antiviral activity of the isopropyl ether compounds **B#24**, **B#59**, **B#57** and the carbamate derivatives **B#43**, **B#60**, and **B#58** was assessed against three different HIV-1 enveloped pseudo typed particles with differing tropism including HXB2, YU2, and 89.6 (Table 1). All six of the azaBINOL compounds showed similar activities across all viral variants. Quinol-type 2′-deoxy-8-azaBINOL ether **B#24** in particular showed low micromolar HIV-1 neutralization against each of the strains tested with activity similar to other antiviral drugs such as abacavir.^34^ Although the naphthol-type 2-deoxy-8-azaBINOL ether **B#59** did show minor antiviral activity at high concentrations, its IC_50_ was greater than the concentrations tested and significantly larger in comparison to the closely related quinol-type ether **B#24**, which has an IC_50_ value of 4 - 8 μM. The significant difference in antiviral activity of these two regioisomers (**B#24** and **B#59**) hints towards a highly specific and selective mode of inhibition. The corresponding carbamate derivatives of deoxy-8-azaBINOLs, **B#43** and **B#60**, both showed moderate antiviral activity against all three viral strains but with smaller selectivity indices. Again, neither of the 2-deoxy-8,8′-diazaBINOL derivatives, **B#57** or **B#58**, showed any antiviral activity at concentrations up to 200 μM.

**Table 1.** HIV-1 pseudo-viral assay determined IC_50_ and CC_50_ for selected azaBINOLs.

| Compound | HIV-IC<sub>50</sub> [μM] |  |  | CC<sub>50</sub> [μM] | Selectivity<br>Index<br>CC<sub>50</sub>/IC<sub>50</sub> |
| --- | --- | --- | --- | --- | --- |
|  | HxB2 | YU2 | 89.6 | TZM-bl |  |
| **B#24** | 6.7 ± 0.9 | 8.9 ± 0.6 | 4.7 ± 1.6 | 68.5 ± 17.1 | 14.6 |
| **B#59** | >100 | >100 | >100 | >100 | --- |
| **B#57** | No effect | No effect | No effect | No effect | --- |
| **B#43** | 61.4 ± 3.3 | 77.6 ± 10.8 | 56.6 ± 0.8 | 95.0 ± 13.1 | 1.7 |
| **B#60** | 12.6 ± 1.6 | 14.0 ± 0.8 | 16.8 ± 3.0 | 69.3 ± 4.9 | 5.5 |
| **B#58** | No effect | No effect | No effect | No effect | --- |

Next we used the EASY-HIT full viral infection assay system^35^ to test all six isopropyl ether and carbamate azaBINOL derivatives against fully-infectious, replication competent HIV-1_LAI_ (Table 2). The quinol-type 2′-deoxy-8-azaBINOL ether **B#24** continued to exhibit low micromolar antiviral activity in the EASY-HIT assay system in accordance with results obtained from the HIVpp assay. Surprisingly, the naphthol-type 2-deoxy-8-azaBINOL ether **B#59** exhibited increased antiviral activity. The deoxy-8-azaBINOL carbamate derivatives **B#43** and **B#60** gave similar high micromolar viral neutralization but also displayed comparable cytotoxicity. The analogous 2-deoxy-8,8′-diazaBINOL compounds, **B#57** and **B#58**, showed no antiviral activity or cytotoxicity at concentrations tested up to 200 μM.

**Table 2.**
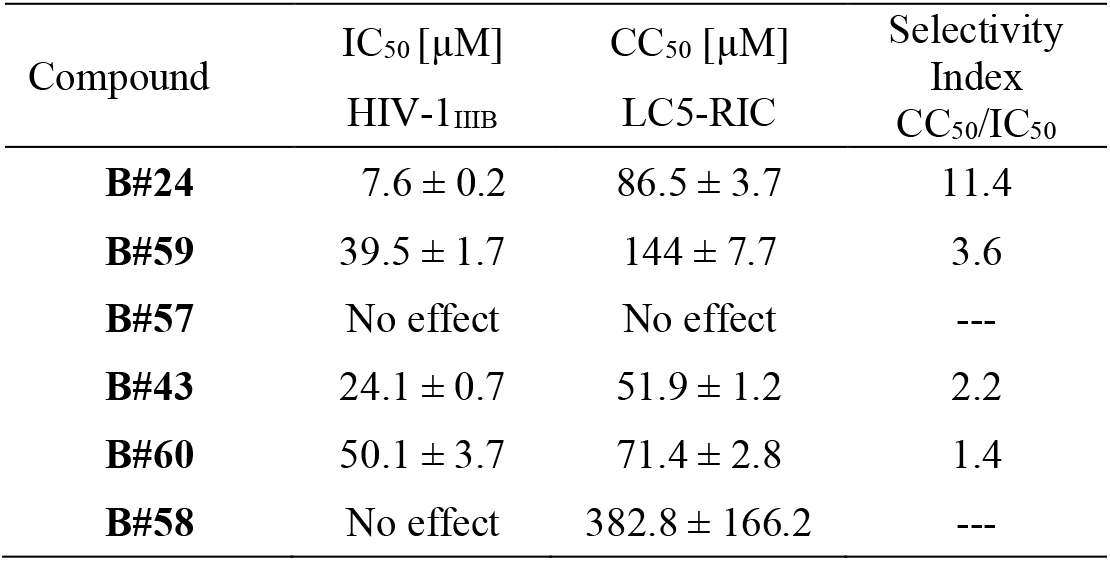
EASY-HIT HIV-1 assay determined IC_50_ and CC_50_ for selected azaBINOLs.

Out of the 41 unique compounds originally screened for anti-HIV-1 activity, one compound, quinol-type 2′-deoxy-8-azaBINOL ether **B#24**, stands out with selective antiviral activity and a favorable selectivity index. Small changes to its structure, including moving the isopropyl ether substituent to the naphthyl ring system (**B#59**), or the introduction of an additional aromatic ring-bound nitrogen atom (**B#57**), results in drastically reduced antiviral activity. While the deoxy-8-azaBINOL carbamate derivatives **B#43** and **B#60** do exhibit antiviral activity in the phenotypic assays, they also show significant cytotoxicity and therefore their apparent antiviral activity is likely due to interference with the cell-based assay. The low micromolar antiviral activity of lead compound **B#24** encouraged us to explore its mode-of-action.

### 3.2 AzaBINOLS are not pan-assay interference compounds (PAINS)

Promiscuous inhibitors in high-throughput screening endeavors often lead to unproductive identification and development of compounds with non-specific activity.^36^ For example, cell-based assays requiring a colorimetric out-read can be inhibited through non-specific mechanisms giving false-positive results.^37^ The cell-based HIVpp used in this study contains a luciferase reporter out read, so we looked to test the capacity of **B#24** to directly inhibit luciferase luminescence in a recombinant luciferase enzyme test (SI Figure 6).^38^ **B#24** showed no inhibition of luciferase activity nor quenching at any concentrations tested verifying that it was not acting on luminescence. Additionally, based on the low solubility and largely hydrophobic surface area of **B#24**, we sought to probe its ability to inhibit HIV-1 through unspecific aggregation effects. We tested the ability for **B#24** to aggregate at higher concentrations in aqueous conditions via an ^1^H-NMR dilution study (SI Figure 7).^39^ Five concentrations of **B#24** were tested from 200 μM to 12 μM in 50 mM phosphate buffer made with D2O and 1% DMSO-d6. No changes were seen in the number of resonances, peak shape, or chemical shift values indicating that **B#24** does not aggregate under the test conditions.

### 3.3 Time-of-addition assay

The single round infectivity assay using HIV-1 enveloped pseudotyped viruses is able to report on inhibition of early stages of infection including cell entry, reverse transcription, and integration steps. The active azaBINOL compound **B#24** inhibited all HIV-1 strains tested regardless of their differing viral tropism (HXB2, YU2, and 89.6). This broad inhibition indicated that the antiviral mode-of-action is unlikely to rely on the viral fusion process, as changing the surface glycoprotein does not affect antiviral activity. Additionally, the EASY-HIT assay indicated that **B#24** is active prior to viral packing and budding. To further delineate the stage of the virus replication cycle inhibited by **B#24**, a time-of-addition assay was performed using HIV-1 pseudotyped particles. Compound **B#24** as well as the standard inhibitors temsavir and efavirenz were added at different time points post exposure of the cell to virus to evaluate their inhibitory activity throughout viral infection (Figure 3). Our results show that **B#24** does not act on the initial viral-entry step when compared to the activity of HIV-1 entry inhibitor temsavir, which loses activity if dosed post viral entry. Instead, **B#24** remains active throughout the assay, but exhibits a subtle decrease in antiviral activity between 6-8 hours post infection. Next, we sought to explore viral enzyme interactions directly to verify if **B#24** interacts with the HIV RT dual functions (DNA polymerase and/or RNase H activity) or HIV integrase.

**Figure 3.**
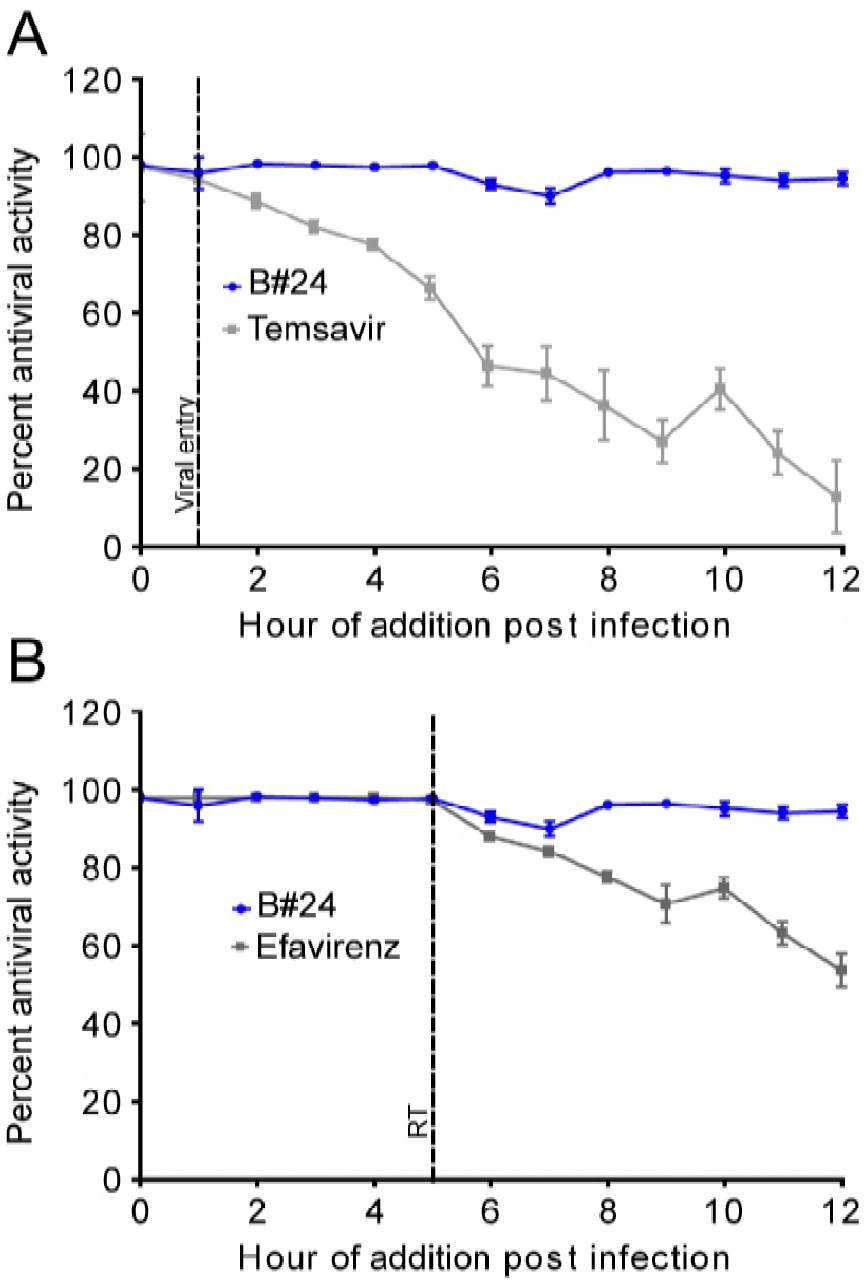
Time-of-addition analysis comparing HIV-1 antiviral activity of **B#24** to antiviral controls at various time points post-infection. TZM-bl cells were infected with HIV-1_YU2_ followed by inoculation with antiviral agents at the indicated time points. A) **B#24** activity profile was compared to an attachment inhibitor (temsavir, 40 nM) and B) a) NNRTI (efavirenz, 40 nM). Results are presented as the mean ± S.D. (*error bars*) of triplicates (n = 3).

### 3.4 Activity of B#24 against HIV-1 reverse transcriptase

NNRTI binding to the HIV RT enzyme occurs at a distant, allosteric binding site and the long-distance effects on the RT polymerase activity are well documented.^10^ In contrast, inhibitors of the HIV RT enzyme that target its RNase H function directly affect the catalytic side with its Mg^2+^ ions, and therefore are often dual inhibitors, with effects on reverse transcriptase and integrase as both require bivalent metals in their active site. We screened **B#24** for inhibitory activity in recombinant protein-based assays to test for HIV-1 RT and/or integrase antiviral inhibition (Table 3). AzaBINOL **B#24** showed no effect against HIV-1 integrase activity in a commercially available kit (ExpressBio, Frederick, MD) at any concentration tested up to 200 μM. When tested against an HIV-1 RT polymerase assay we observed only weak inhibitory effects for **B#24** at concentrations higher than 100 μM, several orders of magnitude weaker than seen in our cell-based assays.

**Table 3.** **B#24** shows specific inhibition against RNase H with an IC_50_ of 14.2 ± 2.8 μM. **B#57** does not show inhibition against any enzyme tested. Data shown represent mean ± S.D. (*error bars*) of triplicates (n = 3).

|  | HIV-1<br>Integrase | HIV-1<br>RT | HIV-1<br>RNase H | *E. coli*<br>RNase H | MuLV<br>RNase H | AMV<br>RNase H |
| --- | --- | --- | --- | --- | --- | --- |
| **B#24** | >100 μM | >100 μM | 14.2 ± 2.8 μM | 44.3 ± 6.4 μM | >100 μM | >100 μM |
| **B#57** | >100 μM | >100 μM | >100 μM | >100 μM | >100 μM | >100 μM |
| Rilpivirine | - | ~1 nM | >10 μM | >10 μM | >10 μM | >10 μM |
| Raltegravir | 1.1 ± 0.2 μM | - | - | - | - | - |

However, since RNase H activity, the second catalytic activity of the HIV-1 reverse transcriptase, is not detected in the above RT-polymerase assay, we investigated the effect of **B#24** on RNase H activity using a previously reported FRET based approach.^40, 41^ Here, we used a pair of oligonucleosides including an 18-mer strand of RNA containing a 3’-fluorescein modification and an 18-mer strand of DNA with a 5’-dabcyl quencher modification. When RNA is cleaved from the RNA/DNA hybrid by RNase H activity, the fluorescent probe is removed from its quenching partner (dabcyl) resulting in fluorescence. We found that **B#24** inhibited RNase H activity of HIV RT with an IC_50_ of 14.2 μM, within the same range as the low micromolar cell-based assay results (Table 3).

Several classes of compounds have shown promising antiviral activity by acting on RNase H activity including N-hydroxyimides,^42^ tropolones,^43, 44^ hydroxypyridonecarboxylic acids,^25, 45, 46^ diketoacids,^47^ vinylogous ureas,^48^ and thienopyrimidinones;^49^ the majority of which target the active site of RNase H through Mg^2+^ ion chelation. With this in mind, we sought to probe whether azaBINOL **B#24** was inhibiting RNase H via the active site using a Mg^2+^ ion chelation and absorbance test. Mg^2+^ ions present in the active site of RNase H are an integral part of its endonuclease function and various inhibitors have been shown to interfere with their chelating properties.^8, 9^ Compound **B#24** was tested for Mg^2+^ binding by assessment of its UV absorption under an increasing concentration of Mg^2+^ following existing procedures.^50^ No UV absorbance changes were observed with the addition of Mg^2+^ up to 120 mM with **B#24** (100 μM) (Figure 4). Therefore, we conclude that **B#24** is not interacting with Mg^2+^ ions and subsequently is not directly inhibiting RNase H activity through active site binding. Instead, it is likely that **B#24** inhibits RNase H enzyme activity allosterically, without affecting the polymerase function of RT. The azaBINOL compound **B#24** adds a new structural class to emerging group of selective HIV RNase H inhibitors including dihydroxy benzoyl naphthyl hydrazone (DHBNH),^24^ various derivatives of vinylogous ureas,^48, 51^ as well as cycloheptathiophene-3-carboxamides (cHTC).^52^, ^53^ Allosteric binding may exhibit fewer side effects compared to active site Mg^2+^ chelating inhibitors and therefore allow for a more favorable therapeutic window.^8, 9^

**Figure 4.**
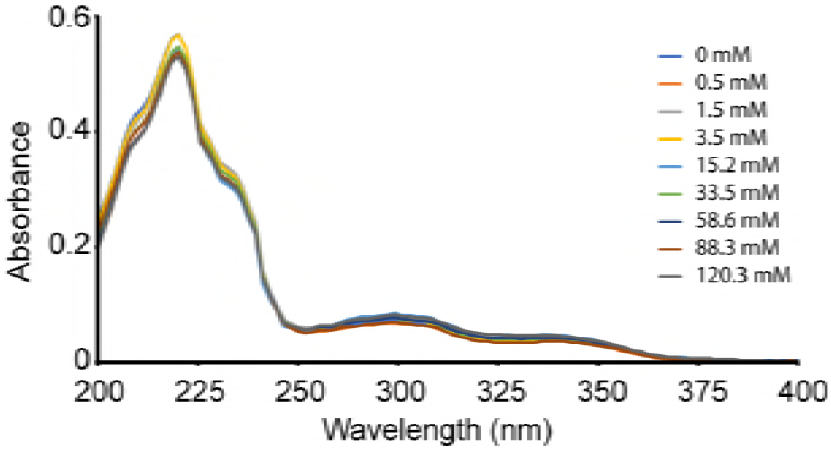
Mg^2+^ ion chelation assay. UV absorbance spectra of **B#24** (100 μM) in the presence of varying concentrations of Mg^2+^ ion. No concentration dependent change of absorbance of **B#24** was seen at any wavelength scanned.

### 3.5 HIV-1 reverse transcriptase binding

To show that **B#24** inhibits RNase H activity by binding the HIV-1 reverse transcriptase enzyme, we explored direct binding to immobilized HIV-1 RT using bio-layer interferometry (BLI). BLI allows for the real time measurement of binding affinities between ligands and analytes of varying size using single-use, fiber optical sensors. BLI measures association (*k*_on_) and dissociation (*k*_off_) rates directly from full spectrum wavelength shifts associated with interference pattern changes derived from binding events at a sensors tip to determine binding affinities (K_D_).^54^ Recombinant-wild type HIV-1 p66/p51 RT (NIH Aids Reagents) was immobilized via amine coupling onto BLI biosensors. **B#24**, selected other azaBINOLs, and control compounds (rilpivirine positive control, raltegravir negative control) were prepared in 1x kinetics buffer at multiple concentrations. Each compound was tested at multiple concentrations, responses were globally fit using a 1:1 binding model, and these fits were used to calculate *K*_D_ values. The binding curves obtained for **B#24** are shown in Figure 5. In good agreement with cell-based antiviral assays, **B#24** shows a concentration-dependent binding to HIV-1 RT. Control compound rilpivirine also showed concentration dependent binding curves while raltegravir, an HIV-1 integrase inhibitor showed no binding at any concentration tested as expected (SI Figure 4). Although the acquired *K*_D_ of 38 μM associated with **B#24** is higher than the IC_50_ in cell-based assays, it should be noted that the affinity value cannot be directly related to neutralization efficacy, as seen in many high-affinity, non-neutralizing HIV antibodies.^55–57^ No binding was observed for the related azaBINOLs of interest from the compound library (SI Figure 7). In summary, the BLI binding data obtained reveals a striking correlation between the binding affinity of **B#24** towards its target HIV RT and its antiviral activity.

**Figure 5.**
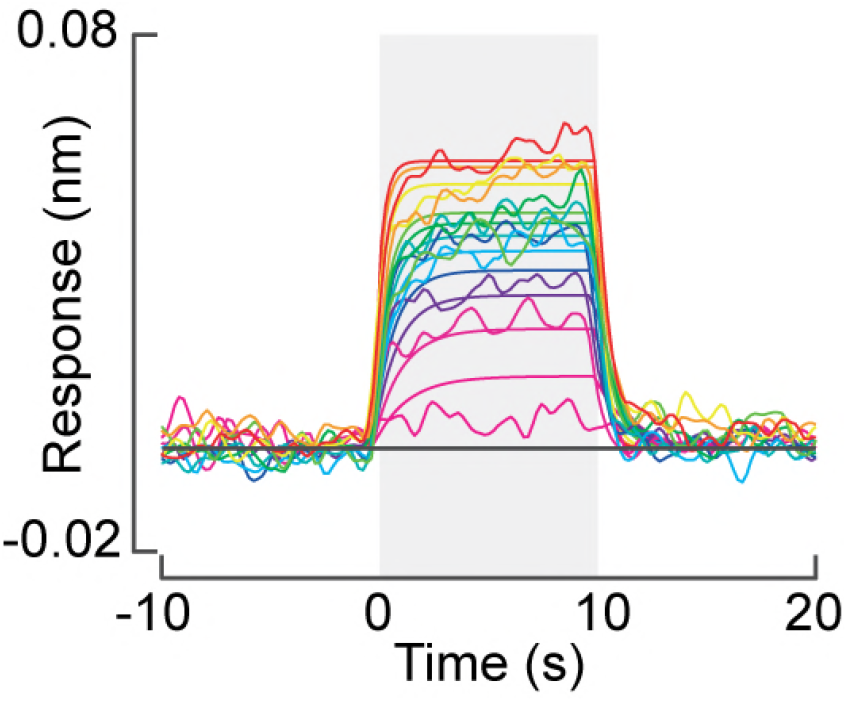
BLI response of inhibitor **B#24** binding to HIV-1 RT enzyme. **B#24** shows a dose-dependent binding to HIV-1 RT with a K_D_ of 38 μM with a 95% confidence interval between 36 - 41 μM. **B#24** was analyzed at 11 concentrations: 10, 20, 30, 40, 50, 60, 70, 80, 120, 160, and 200 μM. Data was fit globally using a 1:1 binding model.

## 4. Discussion and conclusion

RNase H inhibition of HIV-1 RT is an under explored and under-utilized mechanism of inhibition. The azaBINOL compounds reported here represent novel scaffolds that inhibit HIV via an underexplored allosteric mechanism with low toxicity and high specificity. In particular, the isopropyl ether derivative of 2′-deoxy-8-azaBINOL (**B#24**) exhibits with potent and specific antiviral activity against HIV RNase H. We utilized time-of-addition experiments and recombinant enzyme assays to show that **B#24** specifically inhibits RNase H. In addition, we used BLI to show that **B#24** binds to HIV RT via a 1:1 binding mechanism with a binding affinity (K_D_) of 38 μM. Although a number of azaBINOL compounds within the screened library were found to have limited solubility in the cell-based assays as expected, aggregation and nonspecific inhibition was tested thoroughly and can be dismissed for lead compound **B#24** (SI Fig 6,7).

Clinically approved HIV-1 NNRTI’s including nevirapine, efavirenz, and recently rilpivirine, exhibit antiviral activity against HIV-1 by allosterically inhibiting RT through hydrophobic interactions at the NNRTI binding site.^24, 25, 58, 59^ Binding to the NNRTI binding site often results in combined effects in inhibiting DNA polymerase and RNase H activity through conformational shifts in the enzyme.^60^ Only a few privileged structures like the acyl hydrazones,^24^ vinylogous ureas,^48^ cHTC’s,^53^ and now **B#24** have been identified with potent HIV-1 inhibitory effects through specific RNase H inhibition.

We predict that the naphthyl moiety of the azaBINOL compounds may bind to a hydrophobic surface present on the RNase H domain of HIV-1 RT^25^ while leaving the substituted quinoline space to further interact with the protein. In the case of the 2-deoxy-8,8′-diazaBINOL derivatives (**B#57**, **B#58**) or the naphthol-type regioisomer of ether **B#24** (**B#59**), the antiviral activity is absent or significantly reduced likely due to the reduced hydrophobic surface and increased steric interactions, respectively. Ongoing research in our laboratory will explore if **B#24** truly binds allosterically to RNase H and if the binding affinity and HIV-1 neutralization can be enhanced by chemical modifications.

Future studies are planned to optimize the azaBINOL core structure to increase HIV-1 activity by improving HIV-1 binding, solubility, and to reduce cytotoxicity. It is anticipated that the absolute configuration of the biaryl system will influence biological activity but to what extent remains an open question since a majority of the compounds evaluated herein were tested in racemic form only. Work is in progress to determine the configurational stability of compounds such as **B#24** and bioassays of these materials in enantioenriched form will be conducted should their racemization half-lives prove to be high enough for such an effort to be meaningful.

In summary, we discovered that an isopropyl ether derivative of an aza-analog of the archetypal axially chiral biaryl ligand BINOL, inhibits HIV-1 infection in vitro. Based on the identified RNase H inhibition, direct HIV-1 RT binding, and the known hydrophobic binding surfaces associated with allosteric inhibiton,^58^ we believe that **B#24** is a novel allosteric inhibitor of HIV-1 RT-RNase H activity. While biaryl compounds of natural origin have long been known to exhibit significant biological activity,^14, 61^ this study suggests that further investigations of the bioactivity of artificial biaryls, designed purely with synthetic utility in mind, are warranted.

## 5. Experimental Section

### 5.1 General experimental procedures

UV spectra, luminescence and absorbance readings were recorded on a BioTek Synergy HT plate reader. NMR spectra were acquired on Bruker Avance III 400 MHz, Bruker Avance III 500 MHz, Bruker Avance III 700 MHz, and Bruker Ascend 800 MHz spectrometers, equipped with either a 5 mm TXI probe (500 MHz), a 5 mm BBO probe (400 MHz and 500 MHz), or a 5 mm TCI cryoprobe (700 MHz and 800 MHz), with the appropriate solvent signals used as an internal calibration standard [for CDCl_3_: δ_H_ (*C*HCl_3_) = 7.26 ppm, δ_C_ (*C*DCl_3_)= 77.2 ppm]. Numbers in parentheses following carbon atom chemical shifts refer to the number of attached hydrogen atoms as revealed by the DEPT spectral editing technique. Infra-red spectra were recorded on a Perkin Elmer Spectrum II FT-IR using a thin film between NaCl plates. Low (MS) and high resolution (HRMS) mass spectra were obtained using electrospray (ES) ionization on a Waters SYNAPT instrument interfaced with a Shimadzu LC20ad liquid chromatograph. Ion mass/charge (*m/z*) ratios are reported as values in atomic mass units. Preparative chromatographic separations were performed on silica gel 60 (35-75 *μ*m) and reactions were followed by TLC analysis using silica gel 60 plates (2-25 *μ*m) with fluorescent indicator (254 nm) and visualized by UV or phosphomolybdic acid (PMA).

### 5.2 Materials

Non-commercially available BINOL and azaBINOL compounds tested were previously synthesized and characterized at Oregon State University. Preparation details and characterization data for previously undescribed compounds (**B#24**, **B#57**, **B#58**, **B#59**, and **B#60**) are given below. Unless otherwise stated, all solvents were purchased from ThermoFisher Scientific (Waltham, MA) and reagents for chemical synthesis were purchased from Sigma-Aldrich (Milwaukee, WI) and used as received. TZM-bl cells and the HIV-1 inhibitors raltegravir, efavirenz, and rilpivirine were obtained through the NIH Aids Reagent Program.^62–67^ HEK 293T cells were a kind gift from Dr. Pastey Manoj (Oregon State University). HIV-1 viral expression plasmids (pSG3, pHxB2, and pYU2) were obtained as a generous gift from Carole Bewley (NIH, NIDDK). Dulbeco’s modified eagle medium (DMEM) was purchased from VWR. PBS buffer was purchased from Gibco (ThermoFisher Scientific). Trypsin/EDTA (0.25%/2.21 mM) and penicillin/streptomycin solutions were attained from Corning Life Sciences (Corning, NY). Fetal Bovine Serum (FBS) was obtained from Atlanta Biologicals (Flowery Branch, GA). QuantiLum Recombinant Luciferase and the BrightGlo Luciferase Assay System were acquired from Promega (Madison, WI). The HIV-1 inhibitor temsavir was acquired from ViiV Healthcare (Brentford, UK). All compounds received were tested for purity and identity via LCMS analysis before use.

### 5.3 Synthesis of new compounds and characterization data

*Representative procedure for preparation of isopropyl ethers.* 7-(Isopropyloxy)-8-(naphth-1-yl)quinoline (**B#24**): A stirred solution of 7-hydroxy-8-(naphth-1-yl)quinoline (4, 20 mg, 0.074 mmol)^26^ in reagent grade DMF (0.5 mL) at rt under Ar was treated with NaH (12 mg, 60 wt.% in mineral oil, 0.30 mmol). Effervescence was observed. After stirring for 30 min, neat 2-bromopropane (0.030 mL, d = 1.31, 39 mg, 0.317 mmol) was added. The reaction mixture was then heated to 45 °C and stirring continued for 18 h. After this time, the mixture was allowed to cool to rt and partitioned between EtOAc (5 mL) and H_2_O (5 mL). The aqueous phase was extracted with EtOAc (5 mL) and the combined organic phases were washed with H_2_O (5 mL) and brine (5 mL), then dried (Na2SO4) and concentrated *in vacuo.* The residue was purified by column chromatography (SiO2, eluting with 15-30% EtOAc in hexanes) to afford ether **B#24** (20 mg, 0.064 mmol, 86%) as a colorless oil: IR (neat) 2976, 2927, 1610, 1498, 1307, 1259, 1111, 773 cm^−1^; ^1^H NMR (400 MHz, CDCl_3_) δ 8.76 (1H, dd, *J* = 4.2, 1.8 Hz), 8.17 (1H, dd, *J* = 8.2, 1.8 Hz), 7.92 (1H, dm, *J* = 8.3 Hz), 7.90 (2H, d, *J* = 9.0 Hz), 7.61 (1H, dd, *J* = 8.2, 7.0 Hz), 7.47 (1H, d, *J* = 9.0 Hz), 7.45 (1H, dd, *J* = 7.0, 1.2 Hz), 7.42 (1H, ddd, *J* = 8.1, 6.7, 1.4 Hz), 7.31-7.21 (3H, m), 4.42 (1H, septet, *J* = 6.1 Hz), 1.05 (3H, d, *J* = 6.1 Hz), 0.98 (3H, d, *J* = 6.1 Hz) ppm; ^13^C NMR (175 MHz, CDCl_3_) δ 157.6 (0), 150.2 (1), 147.0 (0), 137.9 (0), 133.9 (0), 132.9 (0), 129.1 (1), 128.9 (1), 128.5 (1), 128.3 (1), 126.2 (1), 125.8 (1), 125.6 (1), 125.5 (1), 124.1 (0), 119.0 (1), 118.5 (1), 72.6 (1), 22.3 (3), 22.2 (3) ppm (aromatic C-atom signals not fully resolved, 20 peaks observed for 22 unique C-atoms); MS (ES+) *m/z* 314 (M+H)+; HRMS (ES+) *m/z* 314.1544 (calcd. for C_22_H_20_NO: 314.1545).

**7-(Isopropyloxy)-8,8′-biquinolyl (**B#57**)**: 7-Hydroxy-8,8′-biquinolyl (6, 52 mg, 0.191 mmol)^29^ was converted into isopropyl ether **B#57** (46 mg, 0.146 mmol, 77%) by analogy to the synthesis of **B#24** given above. Data for **B#57**: colorless oil; IR (neat) 2925, 1658, 1596, 1496, 1272, 1112, 1045, 829, 796 cm^−1^; ^1^H NMR (400 MHz, CDCl_3_) δ 8.78 (1H, dd, *J* = 3.9, 1.8 Hz), 8.72 (1H, dd, *J* = 4.2, 1.8 Hz), 8.21 (1H, dd, *J* = 8.3, 1.8 Hz), 8.14 (1H, dd, *J* = 8.2, 1.8 Hz), 7.90 (1H, dd, 7.2, 2.5 Hz), 7.88 (1H, d, *J* = 9.1 Hz), 7.71-7.65 (2H, m), 7.47 (1H, d, *J* = 9.0 Hz), 7.34 (1H, dd, *J* = 8.3, 4.2 Hz), 7.22 (1H, dd, *J* = 8.2, 4.2 Hz), 4.46 (1H, septet, *J* = 6.1 Hz), 1.06 (3H, d, *J* = 6.1 Hz), 0.96 (3H, d, *J* = 6.1 Hz) ppm; ^13^C NMR (100 MHz, CDCl_3_) δ 156.6 (0), 150.7 (1), 149.9 (1), 148.8 (0), 147.8 (0), 136.4 (0), 136.3 (1), 136.0 (1), 132.2 (1), 128.8 (1), 128.6 (0), 127.7 (1), 127.3 (0), 126.3 (1), 124.2 (0), 120.7 (1), 119.0 (1), 118.2 (1), 72.5 (1), 22.5 (3), 22.3 (3) ppm; MS (ES+) *m/z* 315 (M+H)+; HRMS (ES+) *m/z* 315.1502 (calcd. for C_21_H_19_N_2_O: 315.1497).

**2-(Isopropyloxy)-1-(quinol-8-yl)naphthalene (**B#59**)**: 2-Hydroxy-1-(quinol-8-yl)naphthalene (5, 75 mg, 0.276 mmol)^26^ was converted into isopropyl ether **B#59** (66 mg, 0.211 mmol, 76%) by analogy to the synthesis of **B#24** given above. Data for **B#59**: colorless oil; IR (neat) 2976, 2927, 1716, 1593, 1496, 1371, 1235, 798, 750 cm^−1^; ^1^H NMR (400 MHz, CDCl_3_) δ 8.80 (1H, dd, *J* = 4.2, 1.8 Hz), 8.24 (1H, dd, *J* = 8.3, 1.8 Hz), 7.94-7.88 (2H, m), 7.84 (1H, dm, *J* = 8.1 Hz), 7.68-7.64 (2H, m), 7.41 (1H, d, *J* = 9.0 Hz), 7.37 (1H, dd, *J* = 8.3, 4.2 Hz), 7.31 (1H, ddd, *J* = 8.0, 6.6, 1.4 Hz), 7.21 (1H, ddd, *J* = 8.5, 6.6, 1.3 Hz), 7.15 (1H, dm, *J* = 8.5 Hz), 4.36 (1H, septet, *J* = 6.1 Hz), 1.05 (3H, d, *J* = 6.1 Hz), 0.89 (3H, d, *J* = 6.1 Hz) ppm; ^13^C NMR (175 MHz, CDCl_3_) δ 153.5 (0), 150.3 (1), 147.8 (0), 136.9 (0), 136.3 (1), 134.4 (0), 132.5 (1), 129.6 (0), 129.4 (1), 128.6 (0), 128.1 (1), 127.7 (1), 126.3 (1), 126.1 (1), 126.0 (0), 125.9 (1), 123.7 (1), 121.0 (1), 118.5 (1), 72.9 (1), 22.6 (3), 22.4 (3) ppm; MS (ES+) *m/z* 314 (M+H)+; HRMS (ES+) *m/z* 314.1546 (calcd. for C_22_H_20_NO: 314.1545).

*Representative procedure for preparation of carbamates.* **7-[(Diethylamino)carbonyloxy]-8,8′-biquinolyl (B#58)**: A stirred solution of 7-hydroxy-8,8′-biquinolyl (6, 43 mg, 0.158 mmol)^29^ in pyridine (1.0 mL) at rt under Ar was treated with neat diethylcarbamoyl chloride (0.080 mL, d = 1.07, 86 mg, 0.632 mmol). The resulting solution was heated to 100 °C and stirred for 24 h. After this time, the mixture was allowed to cool to rt and partitioned between EtOAc (10 mL), H_2_O (15 mL) and sat. aq. NaHCO_3_ (5 mL). The aqueous phase was extracted with EtOAc (10 mL) and the combined organic phases washed with H_2_O (5 mL) and brine (5 mL), then dried (Na_2_SO_4_) and concentrated *in vacuo.* The residue was purified by column chromatography (SiO_2_, eluting with 3% MeOH in CH_2_Cl_2_) to afford carbamate **B#58** (45 mg, 0.121 mmol, 77%) as a colorless oil: IR (neat) 2930, 1715, 1594, 1417, 1263, 1208, 1157, 796 cm^−1^; ^1^H NMR (400 MHz, CDCl_3_) δ 8.81 (1H, dd, *J* = 4.3, 1.7 Hz), 8.79 (1H, dd, *J* = 4.2, 1.7 Hz), 8.22 (1H, dd, *J* = 6.6, 1.7 Hz), 8.20 (1H, dd, *J* = 6.5, 1.7 Hz), 7.94 (1H, d, *J* = 8.9 Hz), 7.90 (1H, dd, *J* = 8.1, 1.3 Hz), 7.76 (1H, dd, *J* = 7.1, 1.4 Hz), 7.68 (1H, dm, *J* = 7.3 Hz), 7.64 (1H, d, *J* = 8.8 Hz), 7.35 (1H, t, *J* = 4.4 Hz), 7.33 (1H, d, *J* = 4.4 Hz), 3.17-3.06 (2H, m), 2.68 (1H, dq, *J* = 14.2, 7.1 Hz), 2.49 (1H, dq, *J* = 14.2, 7.0 Hz), 0.95 (3H, t, *J* = 7.0 Hz), 0.32 (3H, t, *J* = 7.0 Hz) ppm; ^13^C NMR (100 MHz, CDCl_3_) δ 153.8 (0), 150.7 (1), 150.3 (1), 150.1 (0), 148.4 (0), 147.5 (0), 136.2 (1), 136.2 (1), 135.0 (0), 132.4 (1), 130.4 (0), 128.6 (0), 128.4 (1), 128.1 (1), 126.6 (0), 126.3 (1), 123.4 (1), 120.9 (1), 120.4 (1), 41.9 (2), 41.2 (2), 13.3 (3), 13.2 (3) ppm; MS (ES+) *m/z* 372 (M+H)+; HRMS (ES+) *m/z* 372.1716 (calcd. for C_23_H_21_N_3_O_2_: 372.1712).

**2-[(Diethylamino)oxycarbonyl]-1-(quinol-8-yl)naphthalene (**B#60**)**: 2-Hydroxy-1-(quinol-8-yl)naphthalene (**5**, 50 mg, 0.184 mmol)^26^ was converted into carbamate ****B#60**** (59 mg, 0.159 mmol, 86%) by analogy to the synthesis of **B#58** given above. Data for ****B#60****: colorless oil; IR (neat) 2973, 2931, 1713, 1419, 1269, 1213, 1159, 982, 799, 750 cm^−1^; ^1^H NMR (400 MHz, CDCl_3_) δ 8.83 (1H, dd, *J* = 4.2, 1.8 Hz), 8.23 (1H, dd, *J* = 8.3, 1.8 Hz), 7.96 (1H, d, *J* = 8.9 Hz), 7.92 (1H, dd, *J* = 8.1, 1.5 Hz), 7.91 (1H, dm, *J* = 8.2 Hz), 7.74 (1H, dd, *J* = 7.1, 1.5 Hz), 7.65 (1H, dd, *J* = 8.1, 7.2 Hz), 7.52 (1H, d, *J* = 8.9 Hz), 7.42 (1H, ddd, *J* = 8.1, 6.3, 1.6 Hz), 7.39 (1H, dd, *J* = 8.2, 4.2 Hz), 7.33-7.24 (2H, m), 3.15-3.08 (2H, m), 2.74 (1H, dq, *J* = 13.9, 6.9 Hz), 2.53 (1H, dq, *J* = 14.2, 7.3 Hz), 0.94 (3H, t, *J* = 7.0 Hz), 0.32 (3H, t, *J* = 6.9 Hz) ppm; ^13^C NMR (100 MHz, CDCl_3_) δ 154.2 (0), 150.4 (1), 147.2 (2C, 0), 136.7 (1), 135.4 (0), 134.0 (0), 132.6 (1), 131.8 (0), 129.2 (1), 128.6 (2C, 0), 128.3 (1), 128.1 (1), 126.5 (1), 126.3 (1), 126.2 (1), 125.1 (1), 122.5 (1), 121.2 (1), 42.0 (2), 41.3 (2), 13.3 (2C, 3) ppm; MS (ES+) *m/z* 371 (M+H)_+_; HRMS (ES+) *m/z* 371.1761 (calcd. for C_24_H_23_N_2_O_2_: 371.1760).

### 5.4 Cell culture

TZM-bl and HEK 293T cells were grown in DMEM supplemented with 10% (v/v) FBS, penicillin (100 units/mL), and streptomycin (100 μg/mL). Cells were maintained in a humidified incubator at 37°C with 5% CO_2_. The passage number of cells used in experiments never exceeded 20 passages. All cell lines were tested mycoplasma-negative by real-time PCR (MycoSolutions mycoplasma detection kit, Akron Biotech, Boca Raton, FL).

### 5.5 Pseudovirus production

HIV-1 pseudotyped viruses were prepared as previously described.^68^ Briefly, HEK 293T cells were transfected with an envelope expression plasmid (pHxB2, pYU2, or p89.6) and an envelope deficient HIV-1 backbone vector (pSG3_Δenv_) using XtremeGENE HP DNA Transfection Reagent (Roche). After 24 hours of incubation at 37°C and 5% CO_2_, the growth media was replaced with fresh media followed by an additional 24 hours of incubation. Cellular supernatant was collected and passed through a 0.45 μm filter to give pseudoviral stock. 1 mL pseudoviral aliquots were stored at −80°C until further use in neutralization assays. Viral strength was determined through TCID50 calculations. TZM-bl cells were plated at 9000 cells per well in white 96 well plates (Greiner Bio-One) and incubated at 37 C, 5% CO_2_ for 24 hours. HIV-1 pseudoviruses were added over a two-fold dilution series to wells. At 48 hours-post infection, cells were lysed and BrighGlo luciferase substrate (Promega) added. Luminescence reading were immediately recorded and the TCID50 value calculated as 50% of the maximum light output based on control wells.

### 5.6 HIV-1 pseudovirus single-round infectivity assay

Viral infection rates of HIV-1 pseudoviruses in the presence of inhibitors was measured through HIV-1 pseudoviral tat-induced luciferase production in TZM-bl cells as described previously.^68^ TZM-bl cells were plated at 9000 cells/well into 96 well plates (excluding outer wells to avoid edge effects) followed by overnight incubation at 37°C, 5% CO_2_. Inhibitors and pseudoviral particles at a final concentration of 1x (based on TCID50 measurements) were incubated together for 20 min prior to transfer to adherent TZM-bl cells followed by 48 hours incubation at 37°C, 5% CO_2_. Viral infection was quantified based on luminescent readings taken immediately after the addition of BrighGlo luciferin substrate to infected cells in lysis buffer and relative infectivity rates calculated based on infectious and noninfectious vehicle control wells (1% DMSO). Antiviral IC_50_ values were calculated from compound dilution series ran in triplicate.

### 5.7 HIV Full virus Screening (EASY-HIT)

The EASY-HIT assay^35^ is based on HIV-1 susceptible reporter cells (LC5-RIC) that contain a stably integrated fluorescent reporter gene that is activated upon successful HIV-1 infection and expression of the early viral protein Rev and Tat. Briefly, LC5-RIC cells were seeded into black 96-well plates at a density of 10,000 cells per well 24 hours prior to infection. Compounds stocks dissolved at 20 mM in DMSO were screened at multiple concentrations from 0.1 to 200 μM at a final DMSO concentration of 1% to establish IC_50_ curves. After compound addition, LC5-RIC cells were infected by adding HIV-1 inoculum at an MOI of 0.5 to each well of the plate. Cells were incubated at 37°C, 5% CO_2_ for 48 hours after infection and then measured for reporter expression. Reporter expression was determined by measuring the total fluorescent signal intensity of each well using a fluorescence microplate reader at an excitation filter wavelength of 552 nm and an emission filter wavelength of 596 nm.

### 5.8 Cell viability assays

Cell viability of TZM-bl cells was determined by monitoring mitochondrial reductase activity from the reduction of the tetrazolium salt MTT by metabolically active cells.^69^ TZM-bl cells were plated into 96 well plates (Greiner Bio-One) followed by overnight incubation at 37°C, 5% CO_2_. Compounds were added to wells with a final DMSO concentration of 1% followed by an additional 48 hours incubation. After the designated incubation time, MTT reagent (5mg/mL in 1x PBS) was added to each well to a final concentration of 0.5 mg/mL.

MTT containing plates were incubated for an additional 3 hours after which the media was removed, and the reduced purple formazan product dissolved in 50 μL DMSO. Absorbance was measured at 550 nm. Metabolic activity of vehicle-treated cells (1% DMSO) was defined as 100% cell growth. Cell viability of LC5-RIC cultures exposed to HIV inoculum and test compounds was determined by performing a CellTiter-Blue^®^ cell viability assay (Promega) and monitoring the ability of metabolically active cells to convert the redox dye resazurin into the fluorescent product resorufin. LC5-RIC cells were plated into black 96 well plates (Greiner Bio-One) followed by overnight incubation at 37°C, 5% CO_2_. Compounds stocks dissolved at 20 mM in DMSO were screened at multiple concentrations from 0.1 to 200 μM at a final DMSO concentration of 1% followed by an additional 48 hours incubation. After the designated incubation time, CTB reagent (1:5 in cell culture medium) was added to each well. CTB containing plates were incubated for an additional hour after which fluorescence signal of resazurin was measured using a fluorescence microplate reader at an excitation filter wavelength of 550 nm and an emission filter wavelength of 600 nm.

### 5.9 Time-of-addition assay

To gain a further understanding of the mechanism of action of the antiviral azaBINOL compound **B#24**, a time-of-addition experiment was employed utilizing single-round HIV-1 pseudo particles.^70, 71^ TZM-bl cells were plated into 96 well plates (Greiner Bio-One) followed by overnight incubation at 37°C and 5% CO_2_. After 24 hours, plated TZM-bl cell were infected with HIV-1_YU2_ pseudovirus. **B#24** (60 μM), temsavir (40 nM), efavirenz (40 nM), and raltegravir (1 μM) were added to separate wells at the initial viral inoculation time or at set points postinfection (up to 12 hours post-infection). Cells, virus, and inhibitor were incubated for an additional 48 hours followed by quantification of luciferase production to assess viral infection rates. The antiretroviral drug controls chosen (temsavir, efavirenz, raltegravir) reflect inhibition of HIV-1 at different stages of the viral lifecycle (entry/fusion, reverse transcription, and integration, respectively) allowing for a mechanistic comparison to the unknown antiviral compound **B#24**.

### 5.10 HIV-1 integrase enzyme assay

Inhibition of HIV-1 integrase was assessed using an HIV-1 integrase assay kit (XpressBio Life Science Products) according to the manufacturer’s instructions. IC_50_ values were determined through a two-fold dilution series run in triplicate. The normalized percent integrase inhibition was determined using positive and negative control references with a set amount of vehicle solvent (1% DMSO).

### 5.11 HIV-1 reverse transcriptase enzyme assay

Antiviral azaBINOL **B#24** was assessed for inhibition of HIV-1 reverse transcriptase using a commercially available colorimetric reverse transcriptase assay kit (Roche) according to the manufacturer’s instructions. IC_50_ values were determined through a dilution series run in triplicate. Normalized percentages of inhibitory activity were calculated using positive and negative control wells with a set amount of vehicle solvent (10% DMSO).

### 5.12 Polymerase-Independent RNase H assay

A FRET based assay to assess RNase H inhibition of HIV-1 RT was used as previously described.^40^ In a 100 μL reaction containing 50 mM Tris HCl at pH 7.8, 5.8 M MgCl2, 1 M dithiothreitol (DTT), 80 mM KCl, 2 nM HIV-1 RT (NIH Aids Reagent, cat#3555), and 0.25 μM annealed RNA/DNA hybrid (5’-GAU CUG AGC CUG GGA GCU-Fluorescein-3’; 5’-Dabcyl-AGC TCC CAG GCT CAG ATC-3’; Metabion, Germany) was incubated at 37°C for 1 hour. Enzymatic activity was quenched with the addition of 50 μL of ethylenediaminetetraacetic acid (EDTA; 0.5 M, pH 8.0) and fluorescence read using a Biotek plate reader at 490/528 nm excitation/emission wavelength. Data was analyzed by subtracting the value of a vehicle blank (1% DMSO) and reporting inhibitor as a percentage of the control.

### 5.13 Bio-layer interferometry binding analysis

Binding of compounds to HIV-1 reverse transcriptase was detected and monitored in real time using a FortéBio Octet Red 96 BioLayer Interferometer. Recombinant wildtype HIV-1 RT (NIH Aids Reagent, cat# 3555) was immobilized on amine reactive sensors (AR2G) at 25 μg/mL for 1600 seconds in 50 mM acetate buffer at a pH of 7. Compounds at 20 mM in DMSO were diluted to final concentrations in black 96 well plates (Greiner Bio-One) at a final consistent DMSO concentration of 5% in 1x kinetic buffer (PBS pH 7.4, 0.02% Tween-20, 0.1% albumin, and 0.05% sodium azide, FortéBio). Binding affinity (K_D_) was characterized through the analysis of association and dissociation curves at multiple concentrations. All samples were tested in duplicate. Rilpivirine was used as a positive binding control, raltegravir as a negative control. Effects from non-specific binding were removed using double referencing. Residual baseline drift was calculated by fitting the response during baseline periods to an exponential decay and then subtracting this drift. The resulting response curves were globally fit to a 1:1 Langmuir model.^72^ All fits were performed using a constrained, non-linear least squares minimization (MATLAB R2018a, lsqnonlin function implementing the trust region reflective algorithm). Constraints on parameters were used to guide convergence away from non-physical parameter values, but all constraints were inactive at the converged optimum. A parametric bootstrap analysis was used to compute a 95% confidence interval for the computed K_D_ values, using a normally-distributed error with variance estimated from the sum of squared residuals of the model fit (MATLAB R2018a, 100 iterations). Random noise in response curves was filtered prior to plotting using a smoothing spline (MATLAB R2018a, spaps function).

### 5.14 Bivalent metal binding assay

Testing for complexation of **B#24** with Mg^2+^ ions was carried out following previous protocols with adjustments.^50^ In brief, a 1 M stock solution of MgCl2 and a 1 M solution of **B#24** were prepared in 1:1 ethanol/acetonitrile mixtures. **B#24** was diluted to 100 μM and UV absorbance readings recorded using a BioTek synergy plate reader from 200 – 400 nm. 10 μL additions of a 500 mM Mg^2+^ solution containing 100 μM **B#24** (to keep compound concentration consistent) was added stepwise followed by absorbance readings between each addition. The concentration of Mg^2+^ was raised with each addition from 0.5 mM to 120 mM. Alignment, reference subtraction of blanks, and plotting were done on raw data using Microsoft Excel.

### 5.15 Luminescence inhibition assays

To determine if **B#24** was capable of non-specifically inhibiting luciferase activity in the HIV-1 pseudotyped assay, we implemented a cell-free assay with purified recombinant luciferase (Promega) and luciferin (BrightGlo Luciferase Assay System; Promega) following a previous protocol.^38^ Protein and reagents were prepared and stored according to manufacturer’s instructions. Preliminary experiments established the concentration of luciferin to be used in the assay in order to closely mimic the protein signal in single-round infectivity assay and give maximal sensitivity in luminescence readings. Reactions were carried out in white 96 well-plates (Gibco; ThermoFisher Scientific) by combining compounds with 0.5 μg/mL recombinant luciferase in 1x PBS buffer and 1 mg/mL BSA (VWR). Reactions were initiated by the addition of 30 μL luciferin substrate to wells. Luminescence readings were immediately recorded and normalized using appropriate controls with vehicle solvent (1% DMSO). The compounds Luciferase Inhibitor I (VWR) and raltegravir (NIH Aids Reagent Program) were used as positive and negative controls respectively.

### 5.16 Compound aggregation assessment

We observed precipitation of some azaBINOL compounds at concentrations higher than 200 μM in cell media with 1% DMSO. To assess if the antiviral activity from the active azaBINOL compound **B#24** could be due to non-specific aggregation effects, an ^1^H-NMR assay was implemented to characterize compound behavior in aqueous conditions as described previously.^39^ Briefly, **B#24** was solubilized in DMSO-D6 (MilliporeSigma; Burlington, MA) to a concentration of 20 mM. Serial dilution in 50 mM sodium phosphate at pH 6.8 in 100% D2O (MilliporeSigma; Burlington, MA) yielded five samples ranging in concentration from 200 μM to 12 μM. ^1^H-NMR spectra of each sample was obtained over 64 scans on a Bruker Ascend 800 MHz spectrometer equipped with a 5mm TCI cryoprobe. Data was analyzed and plotted using Bruker TopSpin software.

## Author contributions

All compounds were synthesized and characterized by SB, SMS, and PRB. All cell-based assays were done by RO, GN, and AH. Data analysis was performed by RO, AH, JS, RBW, PRB and SL. The manuscript was written through contributions of all authors and all authors have given approval to the final version of the manuscript.

## Acknowledgments

This work was supported by OSU start-up funds (SL). Envelope plasmids (pHxB2, pYU2, p89.6) were a kind gift from Carole Bewley (NIH, NIDDK). Temsavir was kindly provided by Bristol Myers Squibb. This compound has subsequently been by acquired by ViiV Healthcare. The following reagents were obtained through the NIH AIDS Reagent Program, Division of AIDS, NIAID, NIH: TZM-bl cells (Cat# 8129) from Dr. John C. Kappes, Dr. Xiaoyun Wu and Tranzyme Inc. Raltegravir (Cat # 11680) from Merck & Company, Inc. Efavirenz (Cat # 4624) from the Division of AIDS, NIAID. Rilpivirine (Cat # 12147) from Tibotec Pharmaceuticals, Inc. HIV-1 RT (Cat #3555) from Dr. Stuardt Le Grice and Dr. Jennifer T. Miller. We acknowledge the support of the Oregon State University NMR facility funded in part by the National Institutes of Health, HEI Grant 1S110OD018518, and by the M. J. Murdock Charitable Trust grant #2014162.

## Supplementary data

Supplementary data (additional NMR data and figures illustrating compound library antiviral screening, protein assay results, BLI control data, and assessment of nonspecific inhibition) associated with this article can be found in the online version.

